# SONIC: A Benchmarking Paradigm for Brain-Computer Interfaces

**DOI:** 10.1101/2025.09.30.679683

**Authors:** Sean M. Perkins, Michael Trumpis, Michael E. Reitman, Beata Jarosiewicz, Aashish N. Patel, Adam Weiss, Jacob W. Scott, Kurtis Nishimura, Matthew R. Angle, Shaoyu Qiao, Vikash Gilja

## Abstract

Brain-computer interfaces (BCIs) can restore function for individuals with neurological disorders and have the potential to transform the way people interact with digital systems. However, the development of advanced BCI applications, such as fluent speech synthesis, is dependent on the underlying information transfer capacity of the physical neural interface employed. A significant barrier to progress has been the lack of standardized, application-agnostic methods for benchmarking BCI system performance prior to clinical trials. Here, we introduce SONIC, a novel preclinical benchmarking paradigm designed to evaluate the information transfer rate (ITR) of a BCI system. This paradigm treats the brain and BCI as a noisy communication channel, where information is sent into the brain via precisely controlled sensory stimuli and read out by the neural interface. We implemented this paradigm in an ovine model by presenting rapid sequences of pure tones while recording neural activity from the primary auditory cortex with the Paradromics Connexus^©^ BCI, a fully implanted system utilizing high-density intracortical micro-electrode arrays with wireless power and data transmission. A convolutional neural network was used to decode tones based on neural features. Our results demonstrate an achieved ITR of over 200 bits per second (bps), which is the highest reported BCI ITR to date. For reference, this rate exceeds the linguistic information content of human speech. This ITR is achieved with a total neural interface, filtering, and data aggregation delay of 56 milliseconds. Further analysis demonstrated that ITR remains high (> 100 bps) for the lowest total delay tested (11 ms), supporting the needs of latency-sensitive applications (e.g., direct speech synthesis). This work establishes a new benchmark for BCI performance and demonstrates that the Connexus BCI possesses the bandwidth necessary to support highly advanced applications. This benchmark provides a robust framework for preclinical BCI evaluation, enabling principled system design optimization to accelerate the translation of next-generation neurotechnology.

## 1 Introduction

Brain-computer interfaces (BCIs) offer a direct communication pathway between the brain and external devices to restore or extend natural human abilities. As with any nascent technology, the landscape of BCI applications is shaped by the technology’s current and future capabilities. Today, the technology holds profound therapeutic promise for individuals with a wide range of debilitating conditions. For those with paralysis resulting from amyotrophic lateral sclerosis (ALS), spinal cord injury, or stroke, BCIs have demonstrated the potential to restore communication and control [1–28], offering a renewed connection to the world and a means to regain autonomy. Beyond motor restoration, emerging research suggests a vast therapeutic landscape for BCIs, including applications in neurorehabilitation to promote neural plasticity after stroke or traumatic brain injury, and neuromodulation for treating chronic pain (e.g., [29–31]), mood disorders (e.g., [32–34]), and other neuropsychiatric conditions. Looking further ahead, the potential scope for BCI applications may extend beyond sensory and motor functions to eventually include memory (e.g., [35–37]) and other cognitive processes (e.g., [38–40]). For these long-term applications to become viable, it is essential to develop BCI technology capable of interpreting the subtle, distributed neural patterns that underlie cognition.

The success of ambitious BCI applications depends critically on the performance of the underlying neural interface. To meet the requirements of advanced near-term applications and the field’s long-term goals, BCI systems must be capable of decoding complex, high-dimensional neural states [41–43] with exceptional accuracy and temporal resolution. Lower-bandwidth systems fail to capture finer signal features because of limitations in the number of recording channels, spatial resolution, and/or temporal resolution. These limitations result in such systems effectively averaging over neural patterns during sensing, making their downstream recovery impossible. Therefore, a high-bandwidth neural interface is a foundational requirement to isolate complex neural signals and unlock a broad range of advanced BCI applications.

### 1.1 The imperative for a benchmark

The development of next-generation BCI hardware presents a formidable engineering challenge: unlike the iterative design cycles common in software and semiconductor development, end-to-end testing and optimization of implantable medical devices, like BCIs, necessitate clinical trials. Clinical trials are slow, expensive, and must be performed within a legal and ethical context that is designed to prevent learning by failure. In other words, by construction, clinical trials are not a path to rapid progress through iteration. Here, we propose one such benchmarking test that can enable rapid preclinical iteration in advance of clinical trials, allowing for faster, continuous device improvement without putting human beings at risk.

A common objection to relying on engineering tests, rather than application performance, is that these tests may not align with user needs. This concern extends to any field that relies on benchmark-driven development. In creating benchmarks for BCIs, we can draw lessons from other advanced hardware industries, where establishing standardized benchmarking has served as a catalyst for research and development. Consider the semiconductor industry: similar to BCIs, semiconductors have design cycles dependent upon physical device fabrication, and a single device can enable a broad set of applications. For GPUs, benchmarks have driven decades of innovation; benchmarks like 3DMark for graphics and, more recently, MLPerf for AI provide objective comparisons that enable NVIDIA, AMD, and Google to compete on standardized metrics while advancing the entire field.

The key to designing effective benchmark tests in other industries has been to understand the core platform features that enable high performance across applications, and to test those core features quantitatively and reproducibly. If well designed, these tests can be more effective than subjective user reports, because they offer better controls and interpretability. The evolution of GPUs provides a salient example. A focus on fundamental performance metrics, rather than exclusively on user feedback from the initial gaming market, was instrumental in adapting GPUs for general-purpose computing. This transition proved foundational to the subsequent expansion of modern artificial intelligence, a market that has since grown to exceed that of consumer graphics. Although such metrics do not replace user feedback, a properly designed benchmark serves as a guide for the engineering process, directing development toward systems with the highest potential to succeed across planned and unanticipated applications.

The BCI community has begun establishing its own software evaluation frameworks, such as the Neural Latents Benchmark [44], FALCON [45], and research competitions centered on shared intracortical speech BCI datasets [46, 47]. Although these are vital for refining software-based decoding algorithms, they ultimately operate within the physical constraints imposed by the BCI hardware used to generate the benchmarking datasets. In contrast, we designed a benchmark that is hardware-sensitive, facilitating direct comparison of BCI hardware. It is the BCI hardware that sets the absolute ceiling on performance, as no decoding algorithm can recover information from the brain that the hardware fails to capture in the first place. This motivates the need for a preclinical benchmark focused on the intrinsic capabilities of the BCI hardware itself, which includes sensors that measure physiological signals as well as the recording and control systems that provide these measurements to downstream software-based decoding algorithms.

BCI hardware is designed to measure the state of the brain in real time. Therefore, BCI hardware should be assessed by the speed and accuracy with which it infers relevant brain states. For a motor BCI, those states may correspond to the intention to move a muscle or end effector in a specific way, or could be abstract, such as detection of error in an executed movement. However, motor tasks often have intrinsically low information throughput requirements, and are thus not ideal for benchmarking, as they impose a task-specific ceiling on performance. An ideal testbed involves high task-related throughput and allows for precise, fast, and controlled manipulation of the brain into known states. These known states provide ground truth labels for evaluating the efficacy of decoding. By raising the task-specific ceiling, the measured throughput is no longer bottlenecked by the task, enabling more thorough evaluation of the device. A testbed meeting these criteria can thus provide an application-agnostic metric of BCI system performance.

### 1.2 An information-theoretic framework

Fundamentally, a BCI is an information transfer system: it decodes information about the brain’s state and uses that information to drive external actions. The principles of information theory provide a powerful mathematical framework for analyzing such systems [48]. Within this framework, an entire BCI system, from the neural activity in the brain to the recording hardware, signal processing, and decoded output, can be modeled as a noisy communication channel. By providing structured inputs to that channel (e.g., via sensory stimuli), one can measure the speed and accuracy with which those inputs can be decoded via the BCI. Information transfer rate (ITR), typically measured in bits per second (bps), is a standard metric for evaluating this information transfer process. ITR is computed based on the mutual information between channel inputs and outputs, and accounts for how quickly information is flowing through the channel. BCI systems must be able to decode accurately, but if decoding is too slow (e.g., due to the need to aggregate data over large windows), the BCI will not be able to meet the demands of a wide variety of applications that require low-delay decoding (e.g., cursor control). Individual assessments of speed or accuracy can be confounded by the fact that a trade-off exists between these two factors. ITR is thus a particularly useful metric because it quantifies BCI system performance in a manner that combines speed and accuracy.

### 1.3 Rationale for an acoustic-based preclinical paradigm

To create an effective engineering testbed, it is crucial to isolate the performance of the BCI system from other potentially confounding variables. The measured ITR of a complete BCI system may be limited by any of four factors: (1) the complexity and timing of the information being encoded by the brain, (2) the responsiveness of neurons at the implant site to that information, (3) the properties of the BCI hardware, composed of the physical neural interface, readout, and control electronics, and (4) the efficacy of the signal processing and decoding software downstream of the BCI sensor. Factors 1 and 2 vary across applications and brain regions due to differences in task complexity and the structure, magnitude, and task relevance of neural population activity, all of which can impact measured ITR. The proposed benchmarking paradigm is explicitly designed to standardize Factors 1 and 2 and avoid them bottlenecking ITR, so the resulting ITR becomes a direct and sensitive measure of the intrinsic capabilities of the BCI system itself (Factors 3 and 4). The purpose of the benchmark is to measure the capabilities of BCI hardware (Factor 3). However, reconstructing inputs from measured neural activity inherently requires signal processing and decoding algorithms (Factor 4) that are downstream of BCI hardware. To avoid conflating Factors 3 and 4, the benchmark allows practitioners to design downstream algorithms that are best suited for their BCI hardware. This flexibility is intended to afford effective, equitable benchmarking of a variety of BCI hardware designs.

To meet these design goals, we utilized acoustic stimuli with an ovine preclinical model. The ovine model is commonly used for medical device preclinical testing because of its similar size and weight to humans. Additionally, it offers significant advantages for translational neuroscience applications in particular due to its large, gyrencephalic (folded) brain, which is structurally more analogous to the human brain than many other commonly employed model systems [49–52].

We chose the primary auditory cortex as the implant site. Although many initial BCI applications target the motor cortex, the primary auditory cortex provides an ideal engineering testbed for several reasons. First, it allows for the delivery of precisely controlled input signals: acoustic stimuli can be designed with high temporal and spectral resolution, providing a high degree of control over the information being sent to the brain (Factor 1). Second, the neurophysiological response of the auditory cortex to acoustic stimuli is extensively studied and well-characterized, facilitating the validation of recorded neural signals (Factor 2). Third, the auditory cortex exhibits fast neural dynamics, enabling us to probe the upper limits of a device’s ability to recover brain states with the high accuracy and temporal resolution that are critical for any high-performance BCI platform. Fourth, in the ovine model, the primary auditory cortex is accessible at the brain surface, facilitating testing of a wide range of existing and contemplated BCI device form factors, including intracortical microelectrode arrays, electrocorticography (ECoG) devices, and functional ultrasound (fUS) devices. Finally, an acoustic-stimulus-based testing paradigm does not require substantial behavioral training or auxiliary measurement for awake testing (e.g., visual stimulation paradigms generally require fixation, and motor tasks generally require extensive training and/or precise measurement of movement).

## 2 Standard for Optimizing Neural Interface Capacity (SONIC)

The SONIC benchmark (Figure 1) consists of presenting sequences of tones (*X*) to sheep while recording neural activity from the primary auditory cortex via a neural interface. The measured neural activity is transformed into neural features and passed through a decoding algorithm that generates tone predictions 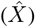. ITR can then be evaluated by comparing *X* to 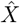.

**Figure 1:**
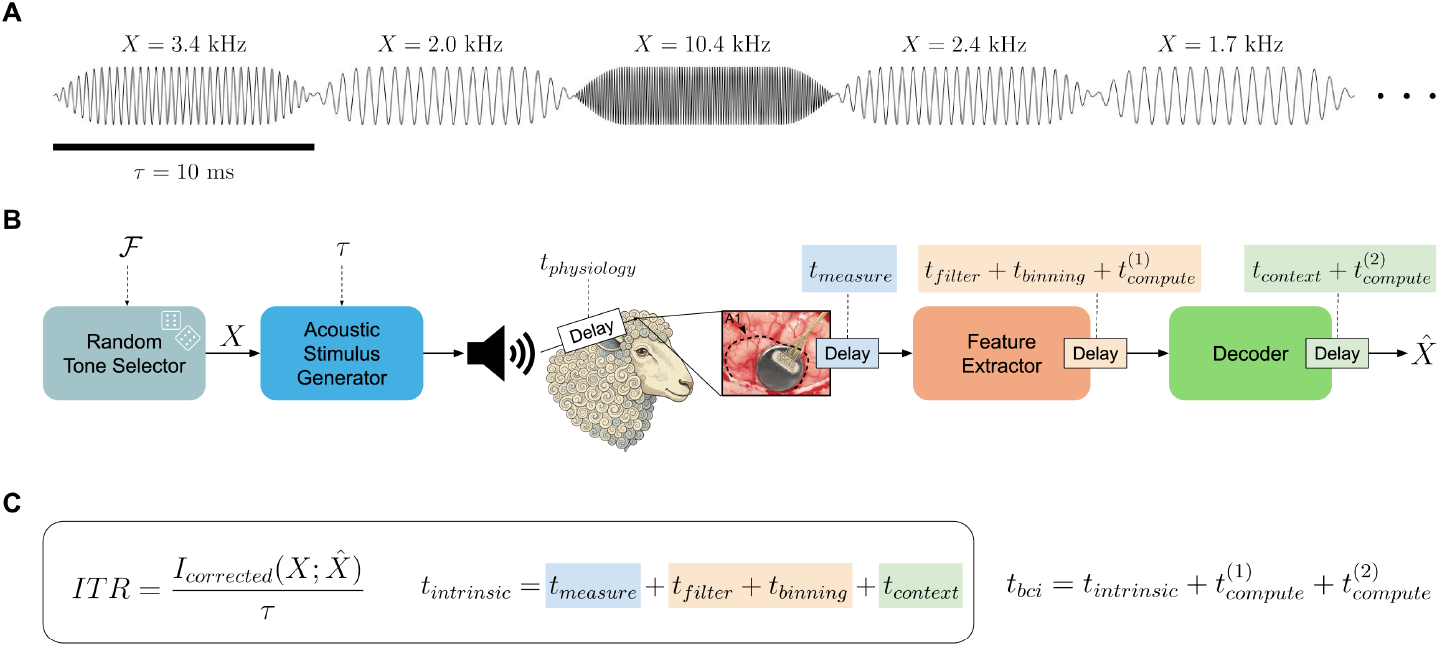
The SONIC benchmark. **A**. A random sequence of pure frequency tones is generated with two free parameters: tone duration (*τ*) and a set of frequencies (ℱ). The frequency of each tone (*X*) is determined by uniformly sampling from ℱ. **B**. Tone sequences are presented to sheep implanted with a neural interface (e.g., Connexus^©^ Cortical Module) in primary auditory cortex (A1, *dashed contour*). Neural activity is recorded, and neural features are extracted and passed through a decoder to generate tone predictions, with each step incurring some delay. **C**. The core output of benchmarking activities is an ITR and an associated intrinsic BCI delay (*t*_*intrinsic*_) that captures delays resulting from measurement via the neural interface, filtering, and data aggregation. The end-to-end delay in an actual BCI system (*t*_*bci*_) also includes delays due to computation time, which are not counted toward the intrinsic BCI delay.

The generation of tone sequences is shaped by two tunable parameters: tone duration (*τ*) and the set of tone frequencies (ℱ) that dictate, via uniform sampling, the sequences of tones (Figure 1A). Tones are presented back-to-back with no silent periods between them. To ensure the choices of *τ* and ℱ do not artificially limit the achieved ITRs, *τ* should be matched to the temporal resolution of the neural interface and ℱ should be matched to the frequency selectivity of neurons at the implant site. For these reasons, *τ* and ℱ are kept as free parameters that should be tuned in exploratory sessions.

ITR reflects both the speed and accuracy of decoding, but is invariant to delays, i.e. there is no penalty for accruing a large delay between neural modulation and a corresponding decoded output. Delayed decoding can impact the viability of BCI applications, so it is important to contextualize a BCI’s ITR with its associated delay (Figure 1B). These delays can come from multiple sources. One such delay is the measurement delay (*t*_*measure*_) between neural activity and its measurement. For devices that measure neurophysiology directly (e.g., intracortical electrodes, ECoG), this delay is based on properties of the BCI sensor (e.g., sampling rate). For devices that measure neurophysiology indirectly, such as functional magnetic resonance imaging (fMRI), functional near-infrared spectroscopy (fNIRS), and fUS, *t*_*measure*_ additionally encompasses the delay between neural activity and corresponding changes in the indirect signal. Additional delays are introduced by downstream signal filtering (*t*_*filter*_), binning for feature extraction (*t*_*binning*_), and accumulation of binned features into a context window by decoding algorithms (*t*_*context*_). Although these operations occur downstream of BCI sensors, they are dependent on the characteristics of the signals captured by the BCI, and specific design choices can impact ITR. We refer to the sum of these delays as *t*_*intrinsic*_, because it encompasses the delays intrinsic to the BCI hardware and downstream algorithmic choices (Figure 1C).

In actual BCI systems, additional delays are present based on the time required for computation (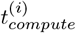 e.g., CPU/GPU runtime), thus the full end-to-end delay in a BCI system (*t*_*bci*_) is given by the sum of *t*_*intrinsic*_ and all computation delays. The precise computation delays are dependent on hardware choices that are not intrinsic to the BCI hardware (and are more readily optimized), therefore, we propose *t*_*intrinsic*_ as the most important delay to report alongside achieved ITRs. The delay between tone presentation and its corresponding neural response (*t*_*physiology*_; i.e., the sensory processing time) is not related to the BCI hardware, and is therefore not relevant to the reported ITRs. The relevant period begins with neural modulation and ends when the BCI produces the corresponding decoded output.

## 3 Results

ITR was evaluated for two sheep with fully-implanted 421-channel wireless Connexus BCI systems across five sessions: one exploratory session followed by four benchmarking sessions. The exploratory session was designed to span a broad range of tone durations (*τ*) and tone frequencies (ℱ), such that these parameters could be subsequently refined for each sheep based on empirical results. The mean achieved ITR across benchmarking sessions for each sheep was 201.9 and 230.1 bps with a *t*_*intrinsic*_ of 56 ms. These represent the highest reported ITRs by a BCI to date.

Refining *τ* is important because neural interfaces capture neural dynamics with varying temporal sensitivity. Selecting the appropriate value for a system is therefore necessary for highlighting system capabilities. Choosing a *τ* that is too short relative to a system’s temporal resolution will underestimate the ITR that could be achieved with a longer *τ*. For very short *τ*, the ovine auditory system itself may struggle to distinguish adjacent stimuli, at which point no neural interface could recover that lost information. However, if *τ* is needlessly long, this limits the number of tones (and thus information) that can be transmitted per unit time. For these reasons, it is important that *τ* remain a free parameter.

Refining ℱ is also essential because tonotopic coverage can vary across sheep and exact implant locations, and ITR will decrease when the stimulus frequencies are mismatched to the frequency selectivity of the recorded neurons. To align with the benchmark’s goal of maximizing neural responsiveness such that measured ITRs reflect the BCI system, not physiological variability, ℱ should be tailored for each implant.

In each exploratory session, we trained a convolutional neural network (CNN) to classify tone frequency from sequences of binned neural features (spike counts and spike-band power) and applied it to held-out data (see Methods). A dense frequency set (ℱ_*initial*_) spanning a broad range was utilized. Model training spanned multiple values for *τ*, but model evaluation occurred separately for each *τ*. Figure 2 illustrates the pattern of errors observed for each sheep at an example tone duration (*τ* = 10 ms).

**Figure 2:**
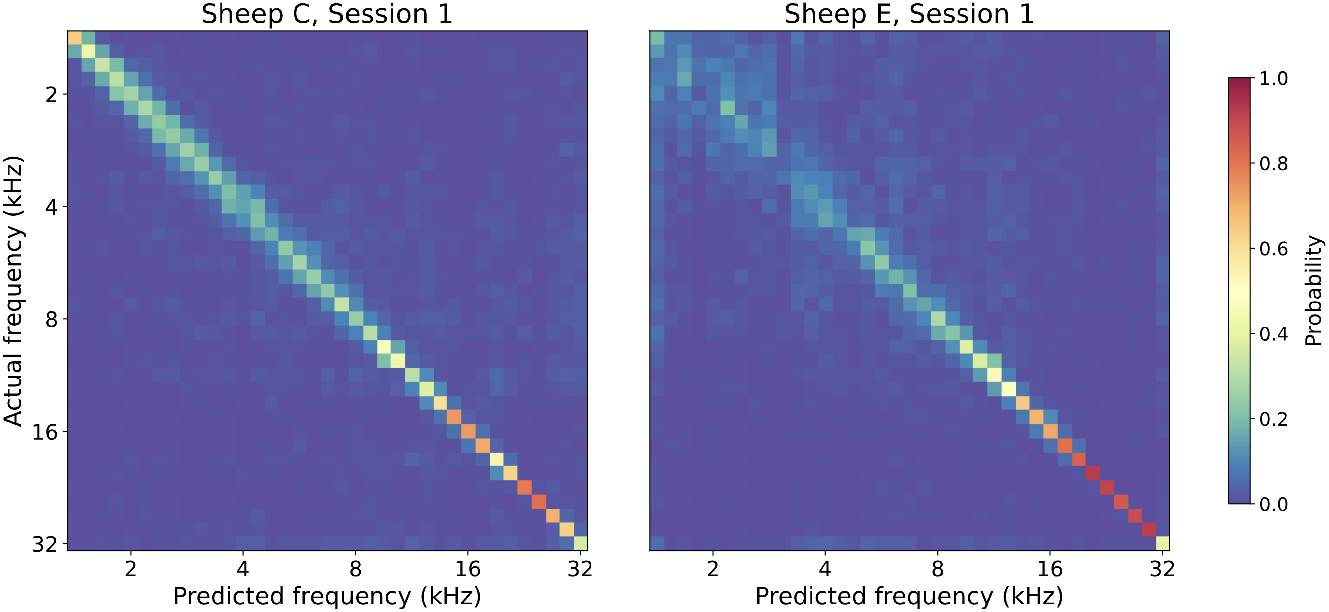
Decoding results in exploratory sessions. In each sheep, an exploratory session was conducted in which sequences of pure tones were presented with frequencies randomly drawn from a large set (ℱ_*initial*_) spanning 1.4 kHz to 32 kHz, and tone durations randomly varied across blocks (4-24 ms, 2 ms increments). Decoders were trained to classify tone frequency from spike counts and spike-band power on data across all tone durations, but were evaluated separately for each duration. The confusion matrices shown here correspond to evaluation on 10-ms tones. Confusion matrix rows are normalized to sum to 1.

Classification accuracy was highest for both sheep at higher tone frequencies (64.7% and 81.9% for frequencies ≥ 17 kHz in Sheep C and E, respectively; accuracy computed using a subset of confusion matrix rows, but retaining all columns). At low frequencies (≤ 4 kHz), Sheep C maintained accuracies well above chance (29.7%, where chance performance is 2.7%), with confusions largely limited to adjacent frequencies, whereas accuracies for Sheep E were much lower (11.9%), with confusions occurring over a broader set of adjacent frequencies. This suggests that lower frequencies are encoded at a coarser resolution in this sheep’s implanted region. These results indicated that ℱ_*initial*_ was suitable for Sheep C, but that a refined set of frequencies would be more suitable for benchmarking sessions with Sheep E. Thus, for subsequent sessions with Sheep E, sparsely spaced frequencies were selected up to 10.5 kHz, and the highest performing frequency range (17-27 kHz) was densely sampled, yielding ℱ_*refined*_. The exploratory sessions also revealed that *τ* = 10 ms led to near-peak ITRs in both sheep (Figure 3). Thus, in benchmarking sessions, model training and evaluation utilized a single *τ*, enabling significantly more training data at this tone duration.

**Figure 3:**
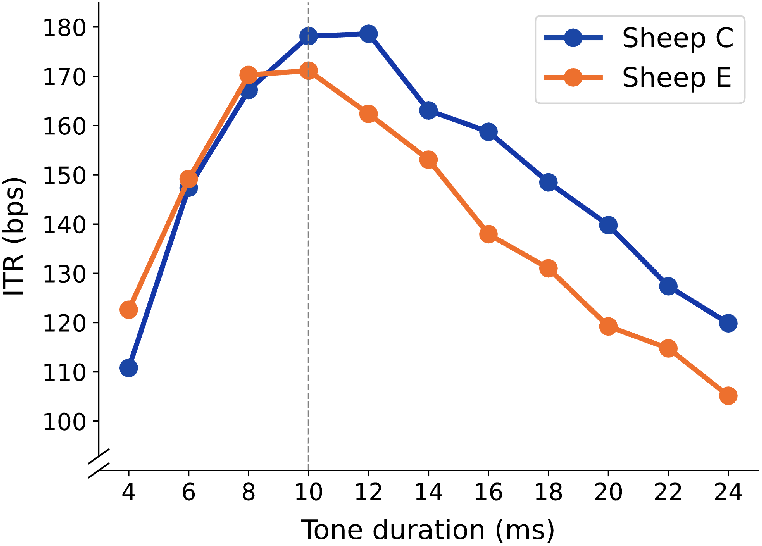
ITR varies with tone duration. ITR was computed separately for each tone duration in the exploratory sessions by training a single decoder across all tone durations and testing it separately on tones of each duration. The 10-ms duration yielded near-peak performance in both sheep, and was the only duration used in subsequent benchmarking sessions.

Benchmarking sessions utilized the same decoding architecture employed in the exploratory sessions, but with the improved choices for ℱ and *τ*. These sessions yielded highly structured confusion matrices (Figure 4), validating these choices.

**Figure 4:**
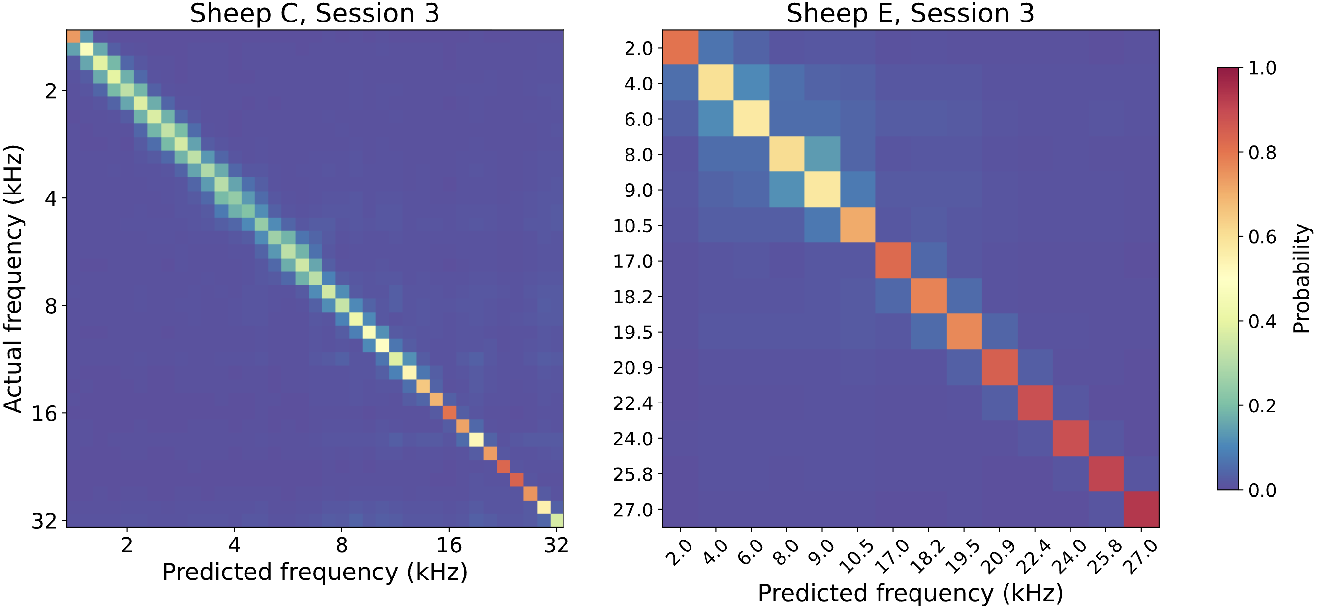
Decoding results in a later session with improved acoustic stimulus parameters. All tone durations for both sheep were 10 ms. The initial tone frequency set (ℱ_*initial*_) from the exploratory session was retained for Sheep C, but Sheep E used a refined set (ℱ_*refined*_) that coarsely sampled the lower frequency range and densely sampled the higher frequency range. Confusion matrix rows are normalized to sum to 1.

We ran four total benchmarking sessions with each sheep using these improved acoustic stimulus settings. The ITRs achieved in each session are listed in Table 1. For both sheep, the mean ITR across sessions exceeded 200 bps. Although the ITRs were similar across the sheep, the accuracies were quite different (45.4% ± 3.1% and 75.9% ± 0.7% in Sheep C and E, respectively; values are reported as mean ± standard deviation, computed across benchmarking sessions). The lower accuracy in Sheep C was expected given how much larger the stimulus set was (37 tones in Sheep C versus 14 in Sheep E). Increasing the number of frequencies in ℱ makes the decoding problem more challenging, decreasing accuracy, but it also increases the average amount of information conveyed by each decoded tone. In fact, the maximum possible ITR that can be achieved by transmitting 10-ms tones is 521 bps when there are 37 possible tones, and 381 bps when there are 14 possible tones. The two sheep thus demonstrate the tradeoff between accuracy and complexity that can occur for any given ITR.

**Table 1.** ITR for benchmarking sessions.

| Subject | Session 2 | Session 3 | Session 4 | Session 5 | Mean ITR (bps) |
| --- | --- | --- | --- | --- | --- |
| Sheep C | 174.6 | 204.0 | 201.6 | 227.2 | 201.9 |
| Sheep E | 228.2 | 231.6 | 225.1 | 235.4 | 230.1 |

The ITRs reported in Table 1 correspond to a *t*_*intrinsic*_ of 56 ms (*t*_*measure*_ < 1 ms, *t*_*filter*_ = 5 ms, *t*_*binning*_ = 5 ms, *t*_*context*_ = 45 ms). The measurement delay is a property of the BCI hardware, while the remaining terms reflect tunable choices. The filter delay is a consequence of zero-phase filtering over a short window of buffered voltages, a viable strategy for real-time decoding that increases spike amplitudes at the expense of a small delay [53]. We selected a filter delay of 5 ms without optimization, but prior work suggests lower delays are practical [53]. The binning and context delays are different from the device and filter delays; the former are shift delays, and the latter are data aggregation delays. Feature binning introduces a delay equivalent to the bin size and decoding algorithms that utilize multiple bins introduce a context delay equivalent to the summed bin sizes (excluding the most recent bin). The ITRs in Table 1 were generated with a data aggregation delay of 50 ms (features binned every 5 ms from 10-60 ms after tone onset).

To understand the sensitivity of the ITRs to data aggregation, we recomputed ITR using all possible data aggregation windows within 0-100 ms after tone onset (Figure 5A). As expected, windows occurring too early or late within the range led to low ITRs, presumably due to misalignment with the most salient neural response to the tones (~ 25-30 ms post-tone-onset, which reflects both *t*_*physiology*_ and *t*_*filter*_). Smaller data aggregation windows also had lower ITRs. However, maximizing over all possible alignments for each data aggregation window (i.e. restricting to data aggregation windows that are centered on the maximal neural response) reveals that ITR remains quite high (> 100 bps) even as the data aggregation window drops down to a single 5 ms bin, which corresponds to a *t*_*intrinsic*_ of 11 ms (Figure 5B). This establishes that the Connexus BCI system retains high information throughput even when operating on brief observation windows, a requirement for low-delay decoding. As discussed below, low delay is an important property for many closed-loop BCI applications (e.g., cursor control or speech synthesis).

**Figure 5:**
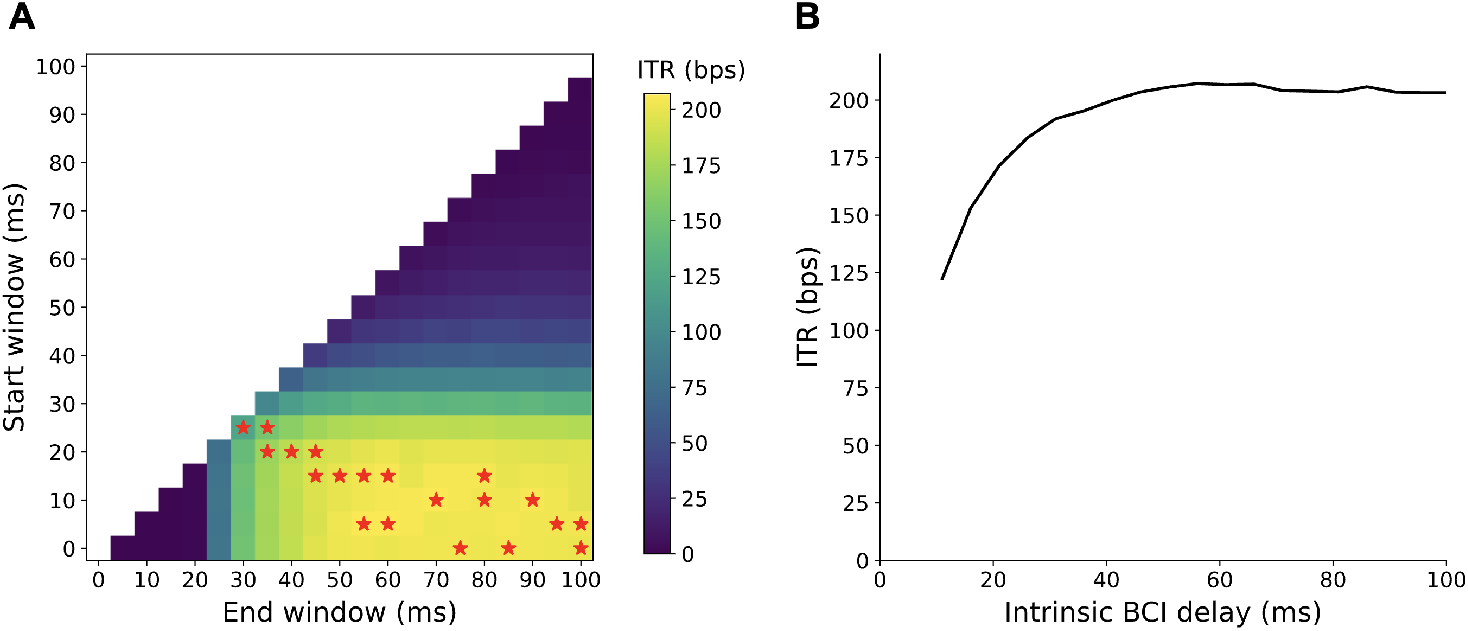
Characterizing ITR across different data aggregation windows. We recomputed ITR for a single session (Sheep E, Session 2) using 5 ms feature bins and varying the data aggregation window over all alignments within 0-100 ms after tone onset (the tone duration was 10 ms). For computational efficiency, only a subset of trials from the session were used and some hyperparameters were changed relative to previous analyses (see Methods). This analysis is only intended to highlight relative differences in ITR across data aggregation windows, but absolute ITRs could likely be further improved (e.g., note in Table 1 that 228.2 bps was achieved for this session using the full dataset and the standard hyperparameters). **A**. ITR is shown for each window in the sweep. *Red stars* indicate the peak ITR over all possible window alignments for a given amount of data aggregation (i.e. the maximum along each diagonal). **B**. ITR is shown as a function of intrinsic BCI delay (*t*_*intrinsic*_), where *t*_*intrinsic*_ is composed of a measurement delay of < 1 ms, a filter delay of 5 ms, and a data aggregation delay that varies across the peak ITRs (*red stars*) from (A).

## 4 Discussion

We designed the SONIC benchmark to measure BCI system information throughput with the goal of isolating the performance evaluation from confounding variables. The choice of an acoustic sensory paradigm in an ovine model reduces the variability inherent in learned motor tasks and bypasses the need for extensive behavioral training. The auditory cortex provides an ideal testbed in which stereotyped, fast neural dynamics can be elicited with an unambiguous ground truth. Among other candidate sensory systems, tactile stimulation may enable faster temporal dynamics, and visual stimulation may enable higher entropy in the stimulus set. However, a major advantage of the auditory system is that the entire perceptual range can be stimulated from a remote source, even without precise localization or directionality. As a result, this benchmarking framework allows for stimulus parameter optimization across a wide range of stimulus space in an untrained animal. Compared to motor tasks, this sensory-based methodology raises the task-imposed ceiling on the measured throughput of a BCI system. Overall, the benchmarking paradigm provides a standardized, reproducible framework to rigorously measure BCI system performance and accelerate the iterative design cycle for next-generation devices prior to resource-intensive clinical trials. In the future, it could also provide a valuable resource for clinicians and potential BCI users who are evaluating the risks and benefits of available technologies.

### 4.1 Information throughput and delay

The applications enabled by a BCI system are dependent on its information throughput and delay characteristics, as well as the tradeoff between these characteristics. As defined in our framework, *t*_*intrinsic*_ encompasses BCI hardware, filtering, and data aggregation delays for feature extraction and decoding. Critically, we demonstrate that the Connexus BCI system achieves an average ITR across sessions above 200 bps in two sheep, with a *t*_*intrinsic*_ of 56 ms. Although the presented analyses were performed offline, evaluation of the Connexus BCI system online confirmed this would correspond to *t*_*bci*_ < 100 ms (exact *t*_*bci*_ is subject to hardware and software choices). When exploring the impact of data aggregation, ITR remained above 100 bps even when *t*_*intrinsic*_ was dropped to 11 ms, indicating available flexibility to reduce delay for applications that require it.

The achieved ITR is not an upper bound for the system, nor is it a guarantee of decoding performance involving other applications and/or brain areas. However, it does demonstrate that the Connexus BCI system is capable of decoding high-complexity information from neural activity with low delay, supporting the potential for highly advanced BCI applications. Each application requires available information throughput to be wielded in a different way. For example, lower information throughput that translates to small mistakes can be tolerated in cursor control (~ 10 bps; [54, 55]), but low delay is essential (e.g., to avoid undesirable cursor dynamics like inertia or orbiting) [56]. In contrast, restoring conversational speech with a lexical output (~ 40 bps) requires high accuracy decoding of language primitives [57], with more modest delay requirements. For reference, the median conversational turn taking delay amongst individuals without speech-related disability is approximately 300 ms [58]. More ambitiously, real-time synthesis of speech (160+ bps based on the speech sound encoder FocalCodec [59]) requires decoding highly complex speech primitives with very low delay. For speech synthesis to match the delays present between motor cortex and production, the delay will need to be less than tens of milliseconds [26]. Achieving a high ITR with low delay in this benchmark can help establish that a device possesses the fundamental characteristics required for a variety of application classes. Conversely, benchmarking a system at a lower ITR can be valuable in appropriately scoping clinical work to lower bandwidth applications.

The tradeoff between throughput and delay is illustrated by recent speech BCI studies, one using surface ECoG grids and two using intracortical microelectrode arrays. Unlike intracortical systems that directly sense spiking activity from a small number of neurons, ECoG-based systems aggregate post-synaptic potentials from a large population of neurons, yielding a signal that is more spatially and temporally smoothed. To compensate for the lower resulting bandwidth, the ECoG BCI employed a bidirectional model that leveraged entire sentence-level context, necessitating a multi-second delay, and limiting the user to a vocabulary of ~ 1,000 words [19]. In contrast, the intracortical BCIs recorded spiking-activity-based features from within the cortex, resulting in higher information content per channel. This higher-resolution signal enabled decoding with much shorter delays, on the order of hundreds of milliseconds, facilitating naturally paced communication with a large vocabulary (> 100,000 words) [17, 21].

Although 200 bps is higher than needed for the applications discussed above, there are practical benefits of exceeding the ITR requirement of a target application. For example, a text-based communication BCI requires an ITR of ~40 bps of intended lexical information to enable communication at rates comparable to speech [57]. A BCI system benchmarked at ~ 40 bps could in theory extract enough information to decode text perfectly at these rates, but in practice it extracts an estimate of neural state that reflects a mixture of application-relevant and application-irrelevant information [60, 61]. Thus, in practice, a system benchmarked at 40 bps would likely receive < 40 bps of application-relevant signal. A system with higher ITR may be more capable of extracting application-relevant information, and may also have capacity to detect additional information. For example, decoding error signals in addition to motor intent can enable a BCI to undo unintended actions [62]. Regions that contain information related to multiple motor actions [63, 64] may enable BCI control of a variety of application types from a single implanted neural interface (e.g., speech and cursor control [24]). Thus, BCI systems with ITRs in excess of individual application requirements may be better equipped to decode a wider variety of behaviors, even if the neural activity associated with those behaviors is not the dominant signal.

### 4.2 The importance of achieved ITR

An important aspect of this benchmark is that the ITRs are achieved, empirical measurements, not theoretical estimates (e.g., channel capacity). A theoretical estimate of ITR is fundamentally a prediction of what could have been achieved had a dataset been collected using a different experiment paradigm. That prediction relies on assumptions, and the computed theoretical ITR can be misleading when those assumptions are violated. For example, the Blahut-Arimoto algorithm [65] utilizes the pattern of errors in a confusion matrix to predict the maximum ITR that could have been achieved had an experiment been conducted with an optimal prior distribution over channel inputs (e.g., tones). However, Blahut-Arimoto assumes the channel is memoryless and will yield an inaccurate estimate if channel outputs depend on previous channel inputs. In many domains, the assumption of memorylessness is reasonable, e.g., an engineered wireless communication system where prior transmitted symbols do not interfere with the current transmission. However, in our benchmarking paradigm, we expect adjacent tones presented in rapid sequence to elicit heavily overlapping neural dynamics. Thus, the assumption of memorylessness is a poor match to the expected data properties. Because of the sensitivity of theoretical ITRs to underlying assumptions, it is important that comparisons in this benchmark rely solely on achieved ITRs.

### 4.3 Limitations and future directions

We recognize that the *t*_*intrinsic*_ associated with benchmarked ITRs may be further reducible via algorithmic optimizations (e.g., using a filter with a smaller *t*_*filter*_). In this sense, *t*_*intrinsic*_ may be considered an upper bound on achievable delays for a neural interface at a given ITR. However, to maintain rigorous benchmarking, such reductions cannot be presumed and should be demonstrated directly via algorithmic updates. For example, the results in Figure 5 motivate investigation into minimizing this upper bound for the Connexus BCI while maintaining a high ITR. This may include reducing filter delay by using a shorter buffer, optimizing bin size (including exploring decoding with a single bin *<* 5 ms), and optimizing the decoding architecture and hyperparameters for operation in the low *t*_*intrinsic*_ regime.

The use of single-frequency tones provides a tractable and reproducible method for establishing this benchmark. However, this stimulus set likely does not probe the full encoding capacity of the neural population in the auditory cortex. Future investigations could employ more complex, feature-rich stimuli (e.g., chords, dynamic spectrograms, and/or naturalistic sounds) to explore the performance limits of BCI systems. One could also explicitly control for fluctuations in attentional state during experiments, which may influence measured ITR.

The results presented in the current study were obtained with fixed parameters for electrode density and cortical coverage. To provide more comprehensive guidance for the ongoing development of BCI hardware, it would be beneficial to undertake a thorough and systematic examination of the tradeoffs that exist between these two factors. It would also be valuable to characterize the relationship between fundamental signal properties (e.g., channel count, sampling rate) and application-specific performance metrics, as well as user feedback. These characterizations can help identify BCI hardware specifications that should be integrated into future benchmarking standards.

We expect the SONIC paradigm to be favorable for devices that average neural activity across space, as primary auditory cortex is known to have a smooth tonotopic map with redundant population coding. Intracortical microelectrode arrays, like the Connexus BCI, are designed to sample spatially local neural activity, which is advantageous for BCI applications involving brain regions with spatially heterogeneous neural responses (e.g., motor cortex). To fully differentiate the capabilities of intracortical arrays from other modalities with less spatial specificity (e.g., ECoG, fUS), it would be beneficial to develop additional benchmarks in cortical areas with more spatially heterogeneous neural activity.

## 5 Conclusion

To accelerate the development of high-performance neurotechnologies, the BCI field requires standardized methods for preclinical evaluation that do not set an artificially low limit on measured performance. The SONIC benchmark presented here provides a robust, application-agnostic paradigm for quantifying the information capacity of any BCI system. By isolating device performance from confounding factors, this approach enables a direct and objective comparison between neural interface technologies. Broader adoption of a benchmark-driven approach to BCI hardware development could move the field more swiftly toward realizing the full therapeutic potential of this transformative technology.

## 6 Methods

### 6.1 Tone presentation

In exploratory sessions (Figures 2-3) for both Sheep C and Sheep E, a set of pure tones were selected in 1/8 octave steps between 1.4 and 32 kHz (ℱ_*initial*_), with 4-24 ms tone durations varied in 2 ms steps. In benchmarking sessions (Figures 4-5, Table 1), all tones had a 10 ms duration. Sheep C continued to use ℱ_*initial*_ in the benchmarking sessions, but Sheep E used a refined set (ℱ_*refined*_), based on decoding performance in its exploratory session. ℱ_*refined*_ consisted of coarsely sampled lower frequencies (2.0 kHz, 4.0 kHz, 6.0 kHz, 8.0 kHz, 9.0 kHz, 10.5 kHz) and densely sampled higher frequencies (17.0 kHz, 18.2 kHz, 19.5 kHz, 20.9 kHz, 22.4 kHz, 24.0 kHz, 25.8 kHz, 27.0 kHz). In all sessions, tone frequencies were randomly selected (uniformly, balanced at the block level) from the above sets and presented consecutively with no inter-tone silent periods (Figure 1). A 1-second silent period occurred between blocks of 8,000-15,000 tones and ≥ 12 blocks were collected in each session. In the sessions using multiple tone durations, each block included every tone duration, tones with the same duration were presented consecutively, and sets of tones with different durations were separated by a 1-second silent period.

### 6.2 Tone generation

Each tone pip consisted of a single sine wave with zero phase at the trial start time, with 2 ms amplitude ramps at the beginning and end to prevent inter-trial clicking. The digital waveform for an entire recording was synthesized in advance and converted to voltages at a 192 kHz digital-to-analog conversion rate using a National Instruments USB-6361 (152805A-03L) Multifunction IO system (NI-DAQ). Acoustic voltages were then amplified with a Tucker Davis Technologies (TDT) SA1 stereo amplifier set to 0 dB attenuation and transduced to sound with a TDT MF1 speaker configured for free-field use, producing sound pressure levels of approximately 80 dB at the distance of the sheep’s ears. Timing was synchronized between the audio production system and the Connexus BCI system via a TimeMachines TM2500C GPS time server providing IEEE 1588 precision time protocol (PTP) and a 10 MHz reference clock. Analog acoustic output and a rising-edge trial time signal were fed directly to the analog input of the NI-DAQ, and the timestamps of the trial-start edges were aligned with broadband electrophysiology signals recorded across all channels from the Connexus BCI.

### 6.3 Signal processing and feature extraction

The extracellular potentials from each of the 421 electrodes, referenced to the Cortical Module case, were amplified and bandpass filtered between ~ 10 Hz and 2.2 kHz through a multi-stage analog hardware pipeline before digitization at a 5.2 kHz sampling rate. Broadband electrophysiology signals were then digitally high-pass filtered (~ 300 Hz, −3 dB cutoff, zero-phase over a rolling 5 ms buffer) and denoised with common average referencing to isolate spike-band activity. Two features were computed from the resulting spike-band activity on each channel: spike counts and spike-band power. Spike counts were obtained by detecting threshold crossings at −3.0 times the root-mean-square (RMS) voltage on each channel, where the RMS was computed from a 2-second period at the start of each recording. Threshold crossings were subject to a minimum inter-spike interval of 1 ms and were counted in 5 ms bins aligned to tone onset for each trial. Spike-band power was computed by squaring each value in the spike-band activity time series and averaging the squared values across each 5 ms bin. For all results excluding the time window sweep presented in Figure 5, the 10 bins spanning the 10-60 ms window following each tone onset were used as input features to the decoder for tone prediction.

### 6.4 Tone decoding

Tone decoding was performed using a two-layer 1D CNN with 400 units per layer, a kernel size of 5 in each layer, and stride of 1 along the time dimension. Each dataset was divided into contiguous segments, and *k*-fold cross-validation was used to generate tone predictions across all trials. The value of *k* varied across datasets (range: 8-16) to ensure that consecutive blocks of trials remained together within a fold, preventing any data leakage across folds due to overlapping feature windows between adjacent trials within a block. During each cross-validation iteration, a test fold and validation fold were held out, and the remaining folds were used for training. Prior to CNN decoding, features were z-scored using normalization parameters learned from a subset of training and validation data. The CNN was trained using a cross-entropy loss function and optimized with AdamW, with a batch size of 64 and utilized a learning rate of 0.001 with a linear decay of 10^−7^ every batch. The early stopping criterion was ITR evaluated on the validation set every 500 batches with a patience of 5. Test set predictions were pooled across folds prior to computing performance metrics to ensure the bias was correctly estimated in ITR calculations.

Minor modifications were made for the analyses in Figure 5 for computational convenience due to the significant number of decoder evaluations necessary. As the purpose of the analysis was to compare data aggregation windows, not to maximize ITR overall, only the first 40% of trials from Sheep E, Session 2 and a single train-validation-test split (75%-12.5%-12.5%) were used. Additionally, the number of convolutional layers was reduced to 1 with a kernel size of 1 (i.e., no convolution). This kernel was necessary for the smallest data aggregation window (a single 5 ms bin) and was therefore applied to all decoding runs in this analysis for consistency. These choices likely result in a lower ITR than application of the methods described above.

### 6.5 ITR calculation

To compute ITR, a confusion matrix was generated based on the presented tones and predicted tones across all trials in a dataset (i.e. predictions were collated across cross-validation test sets). The mutual information (*I*, in bits) between the actual (*X*) and predicted 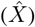 tone frequencies was then computed as:

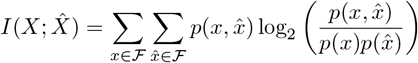

where 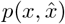 is the joint probability mass function for the random variables *X* and 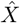, and *p*(*x*) and 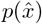 are the associated marginal distributions. This computation is possible because 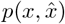 can be generated by normalizing the confusion matrix. In the limit of infinite trials, this estimate of the probability distribution would approach the ground-truth probability distribution and a naive computation of mutual information would be sufficient. However, the mutual information is biased upward when provided a limited trial count. Given there is a non-zero probability of confusion between any two frequency pairs (a reasonable assumption for classification tasks), a first-order approximation of the bias (*b*, in bits) can be computed as described in [66]:

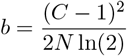

where *C* is the number of classes and *N* is the total number of trials. Increasing the class count increases the bias because it becomes less likely that the outcomes (*C* × *C*) will be adequately sampled by a finite trial count. Bias-corrected mutual information per tone can be computed as

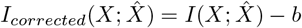

Because the tones are played back-to-back with no inter-tone interval, ITR can be computed in bits per second by normalizing the bias-corrected mutual information by the tone duration (*τ*, in seconds):

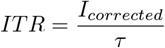

### 6.6 Animal preparation

Two castrated male Suffolk Cross sheep (*Ovis aries*) participated in this study (Sheep C: 76.5 kg; Sheep E: 66.0 kg at the time of implantation). All surgical and experimental procedures complied with the National Institutes of Health *Guide for the Care and Use of Laboratory Animals* and were approved by the Institutional Animal Care and Use Committee (IACUC) of the animal facility. Animals were housed indoors in pens at the animal facility under a 12 hour light/12 hour dark cycle.

Both sheep were chronically implanted with a fully implantable Connexus BCI system comprising a Cortical Module, an Internal Transceiver (IT), and an Extension Lead. The Cortical Module was implanted subdurally within the left primary auditory cortex, and the IT was implanted subcutaneously above the left rib cage, with the Extension Lead tunneled subcutaneously to connect the two components. All neural recordings were conducted during the light cycle while the animals were awake and alert, secured in a stanchion within an electrophysiology rig.

Sheep C and E were part of a cohort for a prospective study evaluating *in vivo* safety and reliability of the Connexus BCI. During Cortical Module implantation, each implant procedure provided different surgical access due to inherent differences in the anatomy and location of the primary auditory cortex in each sheep, resulting in variability in the size and geometry of the implantation target exposed within the craniotomy window. These two sheep were selected for benchmarking activities because they had the largest proportion of electrodes (~ 334 in Sheep C and ~ 341 in Sheep E) implanted in the primary auditory cortex, based on visual identification of pre-implantation anatomical landmarks. This criterion was chosen to maximize coverage of neural populations that encode auditory information.

### 6.7 Connexus BCI device details

The Cortical Module (10 mm in diameter) consists of a 421-channel Platinum-Iridium (Pt-Ir) micro-electrode array (7 mm in diameter), a sealed titanium alloy case which houses an application-specific integrated circuit (ASIC), and an Integrated Lead. The solid Pt-Ir microelectrode array is bonded to a hermetic metal-ceramic feedthrough with an array of 421 vias. The feedthrough vias are bonded to the ASIC on the opposite side, electrically connecting each electrode to the ASIC. Each electrode is 1.5 mm long, 40 µm in diameter, and spaced 300 µm from neighboring electrodes. The Pt-Ir microelectrode array is coated with Parylene-C and the electrode tips are deinsulated for neural recording. The ASIC consists of an array of low-noise amplifiers, filters, and an analog-to-digital converter. The 421 input channels are multiplexed into an 8-wire Integrated Lead, which connects to the Extension Lead and subsequently to the IT. The Cortical Module provides a tunable sampling rate and is capable of acquiring both extracellular action potentials and local field potentials.

The implanted IT receives inductive power from the External Transceiver (ET). The IT distributes power to the Cortical Module and transmits neural data wirelessly through the skin to the ET via a secure near-infrared optical link, supporting speeds of up to 100 Mbps. The ET is subsequently connected to the Control Unit for data acquisition and device configuration. Power delivery and data transmission occur when an ET-IT alignment is established, enabling high-bandwidth neural data transmission without any transcutaneous connections.

## 7 Acknowledgments

We would like to thank the current and former Paradromics employees for their contributions to the design, manufacturing, and testing of the Connexus BCI. We would like to thank our interns—Surya Pandiaraju, Tereza Okálová, Nathan Liu, and Yusuf Ali—for their helpful technical discussions. We would also like to extend our gratitude to the veterinary staff at the animal facility for their support with animal preparation and care.

## 8 Author contributions

Conceptualization: SMP, MT, KN, MRA, SQ, VG

Implantation Surgery: SQ

Experiment Design: SMP, MT, MER, BJ, ANP, SQ, VG

Data Collection: MT, MER, JWS

Data Analysis: SMP, MT, MER, ANP, AW, SQ, VG

Writing: SMP, MT, MER, BJ, ANP, MRA, SQ, VG

Supervision: SQ, VG

